# A multitask neural network trained on embeddings from ESMFold can accurately rank order clinical outcomes for different cystic fibrosis mutations

**DOI:** 10.1101/2023.10.26.564274

**Authors:** Erik Drysdale

**Affiliations:** The Hospital for Sick Children

**Keywords:** Machine Learning, Protein Folding, Cystic Fibrosis

## Abstract

Advancements in AI-based protein structure prediction have opened new research avenues in computational biology. Previous work has shown mixed results when the predicted 3D structure or model embeddings have been used as features for predicting phenotypes. Many existing tools which predict the pathogenicity of variants (e.g. SIFT 4G, Polyphen-2, REVEL, LYRUS) have incorporated structural protein features, but have shown mixed success when applied to mutations on the CFTR gene. This paper explores whether embeddings from the ESM-Fold model, which predicts the 3D structure of a protein from an amino acid sequence, can be used to predict clinical outcomes for different Cystic Fibrosis (CF) genotypes found in the CFTR2 database. A neural network model trained on these embeddings is able to obtain both non-trivial and statistically significant correlations on CF-related phenotypes (sweat chloride, pancreatic insufficiency, and infection rates). Overall, AI-based protein structure prediction models show a promising ability to assess the relative severity of CFTR-based mutations on key phenotypic outcomes associated with CF. The entire processing and analysis pipeline for this work can be found at this [repo](https://github.com/ErikinBC/cftr2_esmfold).

## Introduction

Cystic fibrosis (CF) is a rare autosomal recessive genetic disease that can result in a progressive deterioration of organ function, most notably in the lungs and pancreas (1). Patients with severe CF symptoms can have trouble breathing due to thick and sticky mucus in their lungs (2). Most CF patients that die of their disease do so because of end-stage obstructive lung disease (90%) (3). The Caucasian population is most at risk of CF, although other groups are still at risk (and likely at a higher rate the official numbers would suggest due to under reporting (4)).

The disease’s etiology is caused by mutations to the cystic fibrosis transmembrane conductance regulator (CFTR) gene which causes an alteration in the chloride and bicarbonate transport channel regulated by cyclic adenosine monophosphate (cAMP) (5). This can lead to a pathological build up of mucus in the organs, with the most deleterious impact being on the lungs and the pancreas. As of March 2023, a total of 2114 mutations have been discovered on the CFTR gene (6), with 401 of those being identified as CF-causing (7). The most common mutation is F508del (aka c.1521_1523delCTT, aka p.Phe508del), which is present in 82% of the CF population (8).

Mutations to the exonic (protein-coding) region of the CFTR gene can have disease-causing properties because they can alter the amino acid sequence of the gene. For example, patients with the F508del mutation are missing three DNA codons which code for the phenylalanine amino acid. Even though the other 1479 amino acids in the gene match the wild-type protein, the absence of this critical amino acid impacts the ability of the different CFTR protein domains to take up their proper position (9). As F508del reveals, knowing the physical shape of the mutant protein helps to explain its key pathologies.

Being able to predict the final f olded s hape o f a protein *a priori* was a largely unsolved problem before “transformational” AI-based breakthroughs that began in 2018 (10). Experimental-level accuracy was first d emonstrated b y AlphaFold2 in 2020 for certain types of proteins (11). AlphaFold’s approach has been replicated by other researchers (12), as well as for different architectural approaches like ESMFold or RoseTTAFold (13, 14). ESMFold leveraged techniques from natural language processing including the use of transformers and self-supervised learning on masked sequences of amino acids to pretrain the model before a downstream tuning on the protein folding prediction problem. Unlike other approaches, ESMFold can be used without referencing amino acid sequences of other organisms (known as multiple sequence alignment) and is therefore faster at inference time.

Despite claims that the protein folding problem has been “solved” (15), there is mixed evidence as to whether these tools are sensitive enough to detect small local effects of single mutation, since the change caused may be relatively limited compared to the natural movement of the protein. At least two studies have shown that AlphaFold’s output has a limited correlation when for predicting the impact of mutations relative to the wild-type (16, 17). And while and other papers find different results (18, 19), it is still an open question as to circumstances and domains in which these models can provide insight into the effects of these mutations.

There are a substantial number of bioinformatics tools for predicting the pathogenicity of mutations such as SIFT 4G (20), Polyphen-2 (21), PROVEAN (22), and FATHMM (23).

These tools usually fall into one of three categories: i) sequence based tools, ii) structure based tools, or iii) dynamics based tools. There is an equally large literature on how to stack these tools to improve performance (24, 25).

As most pathogenic mutations appear to affect protein stability (26), protein folding models may be able to produce useful signals for predicting disease severity. Even in the pre-AlphaFold era, classifiers that took into account 3D structures showed some success (21, 27). Other research has shown that an ML model trained on AlphaFold features (referred to as AlphaScore in the paper) improved the accuracy of pathogenicity predictions when combined with other tools (28). However this finding has not been universal (16, 17). The use of existing tools for predicting the pathogenicity of missense mutations on the CFTR gene has been mixed, with one paper showing relatively high sensitivity and specificity (29), whilst another found less accurate results when using a larger sample of CFTR mutations (30). Earlier work showed that CFTR-specific phenotype-optimized sequence ensembles (POSEs) could outperform off-the-shelf sequence based tools (31).

Only one paper in the CF literature, as far as this author can tell, explicitly correlated mutational pathogenicity generated from a model to CF-specific clinical outcomes (32). Thirtyfive different features ranging from existing bioinformatics tools to CFTR-specific protein information were combined and the stacked model showed that the predicted pathogenicity probability was correlated to both chloride conductance and sweat chloride levels on proprietary data (see Figure 4 in their paper) (33). Only patients heterozygous for F508del mutation were included.

Another related research paper showed that two AI-based protein structure models (FoldX and Rosetta) could be used to predict the domain stability on NBD1 and NBD2 of the CFTR gene relative to experimental results (19). However the paper looked at a small number of mutations (59) and did not link this to any clinical outcomes. Like the previously mentioned paper (32), this paper’s analysis correlates a predicted measure to a level of clinical severity, but uses a deep learning based structural model to obtain mutation-based features.

The approach taken by this paper stands out from the existing literature in three ways. First, it considers multiple clinical outcomes types (e.g. not only sweat chloride) and label constructions (e.g. not only F508del heterozygotes). Second, the model is fine-tuned to make predictions directly on CF-specific clinical outcomes rather than binary/ordinal pathogenicity categorizations. Third, by using the embeddings from the ESMFold model, the ML model is able to make predictions for multiple mutation types including insertions, deletions, and duplications, rather than just missense mutations as is done in most of the literature.

## Methods

Note, the entire processing and analysis pipeline for this work can be found at this [repo](https://github.com/ErikinBC/cftr2_esmfold).

### Clinical and genetic data

Average clinical outcomes for four phenotypes (sweat chloride, lung function, pancreatic insufficiency, and infection rates) were obtained from the CFTR2 website for 417 single-variant mutations and 1885 two-variant mutations. Not all variants had clinical information due to small cell censoring – any phenotype with 5 or fewer measurements was censored (see Figure S2). There is substantial variation in (average) clinical outcomes: between mutations (Figure S4), within mutations (Figure S5), between categories (Figure S6), and within categories (Figure S7).

Each variant was linked to an NCBI page and then a consistent genomic location was extracted using the NCBI Variation Viewer for GRCh38. Figure 1 shows the location of 341 exonic mutations found in the CFTR2 database that had an established NCBI page and location. Most mutations are point substitutions (70%), followed by deletions (20%), duplications (8%), and insertions (2%).

**Fig. 1.**
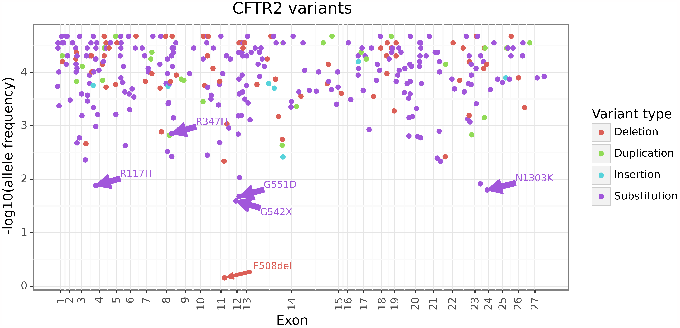
CFTR2 exonic mutations

Using the wild-type CFTR gene, mutations were applied to the reference genome and a new amino acid sequence was calculated for each mutant CFTR gene. Figure S8 shows the empirical CDF of the amino acid length for each version of mutated CFTR gene up to the first stop codon. The wildtype CFTR gene has 1480 amino acids, and around half of the exonic CFTR mutations are (approximately) this length. Mutant CFTR genes can have significantly fewer amino acids than the wild-type due to mutations which add a stop codon. Supplementary Section A provides additional information on how the the data was collected.

### ESMFold

An A100 on a Lambda Lab GPU Cloud was used to perform inference and Supplementary Section B gives further details on the hyperparameter setup. A total of 311 amino acid sequences were passed through ESMFold model. This number represents all exonic mutations that: i) were nonsynonymous, ii) did not have a duplicative amino acid sequence to another mutation, and iii) had an amino acid length of at least 100 when measured to the first stop codon.

Inference using the ESMFold model has a runtime of *O*(*n*^3^), and Figure S9 shows that it takes 4 minutes for an amino acid sequence length of 1500 for a single recycle. From the forward pass, the following embeddings were extracted^1^:

1. s_s - (1, n_amino, 1024): Per-residue embeddings derived by concatenating the hidden states of each layer of the ESM-2 LM stem.
2. s_z - (1, n_amino, n_amino, 128): Pairwise residue embeddings.
3. states - (8,1,n_amino,384): Hidden states from the protein folding trunk.

While the 3D coordinates could also have been used, the embeddings were chosen since ML models are more amenable to a matrix of features which can be reduced to a fixed size. In contrast, had the 3D coordinates had of been used, more complicated algorithms like a 3D-CNN might have needed to be employed to extract features.

Two approaches were used to generate fixed dimensions for the extracted ESMFold embeddings:

1. The average, min, max, and standard deviation of all dimensions from each embedding are taken except the last.
2. The average, min, max, and standard deviation of the cosine similarities between the wild-type and mutant CFTR embeddings are calculated.

This produced a concatenated vector of length 23904, with the first 6144 dimensions coming from approach #1 and the remaining 17760 coming from approach #2. See Supplementary Section B for more information on the feature engineering.

### Phenotypes

A total of four clinical measurements (phenotypes) were obtained for each genotype combination (single- or two-variant), although as noted before some measurements are missing:

1. sweat chloride measurement (mEq/L)
2. lung function (FEV1% = Forced Expiratory) for three age ranges (<10, 10-20, and 20+)
3. % of patients with pancreatic insufficiency
4. % of patients who have had a Pseudomonas infection

Figure S6 shows that many of the phenotype categories are correlated with each other (-37% to +63%). For example, mutations which tend to have higher pancreatic insufficiency will also have higher sweat chloride levels but lower lung function.

The CFTR2 database provides a min-max range of lung function for three different age range (<10, 10-20, 20+). The min, max, and midpoint was also determined for each of these each ranges. An “All-ages” category was also calculated: where the min was the minimum, the max the maximum, and the the midpoint the average of the three age-base categories. A total of 12 lung measurements (the combination of four age categories and three statistics) was therefore available. When combined with the three other clinical measures (sweat chloride, pancreatic insufficiency, and pseudomonas infection rate) there were up to 15 categories of phenotype. While each mutation had up to 15 phenotype measurements, there were different measures for variations of that genotype. For example, the mutation R560T has five different CFTR2 results: i) R560T, ii) R560T-F508del, iii) R560TR560T, iv) R560T-G551D, and v) R560T-N1303K. It is not obvious which is the right phenotype measurement to associate with the R560T genotype. Four label types were therefore used calculate for each mutation:

1. A single-mutation average which includes all patients who had one or more of the alleles (e.g. R560T)
2. The heterozygous combination with F508del (e.g. R560T-F508del)
3. The homozygous combination (e.g. R560T-R560T)
4. The average of all heterozygous combinations (e.g. the average of R560T-F508del, R560T-G551D, R560T-N1303K)

Figure S7 shows the within-category correlation across the four label types, which ranges from an average of 66% to 95%. The heterozygous average (approach #4) has the highest correlation with F508del-heterozygous (approach #2) for the simple reason that of the 219 mutations that have at least one heterozygous measurement for at least one phenotype, 176 only have F508del (80%). In contrast, the homozygous measurements (approach #3) have the lowest correlation, although this depends on the category (see Figure S10). The relationship between the allele-specific average (approach #1) and the heterozygotes (approach #2 & #4) is strong on average (see Figure S11, although there meaningful differences for many mutations). Figure S5 shows why coming up with “one” phenotype for a mutation is challenging since different heterozygotic pairs can have tremendous variation in clinical outcomes. In other words, how bad a mutation is really going to depend on which other mutation it is paired with.

#### Amino Acid Length Adjustment

Mutations whose CFTR gene was about the same length as the wild-type (1480) had better clinical outcomes, on average, than those mutations with stop codons inserted early, particularly for sweat chloride and pancreatic insufficiency. Up to 43% of the clinical variation can be explained by simply knowing the length of the polypeptide chain to the first stop codon (see Figure S12). To prevent the ML model from learning to predict this confounder, a leave-one-out Nadarya-Watson (NW) kernel regression method to get the “length-adjusted” clinical phenotype value. Because the homozygous label for the infection rate category had an R-squared that was less than zero, the label intercept was used instead of the NW estimator.

Preliminary explorations showed that there was limited signal between any of the lung function categories/labels and the ESMFold embeding. Therefore, in the final analysis only 12 labels were used, which were a combination of three categories (sweat chloride, pancreatic insufficiency, and infection rate) and the four label types.

### Algorithm

A simple two-layer feed forward neural network model was used to predict all 12 outcomes simultaneously. The algorithm, hereafter referred to as the “NNet” or “ESMFold-NNet”, had a shared multitask architecture, with each output node being a linear combination of the second hidden layer’s nodes. The NNet used a learning rate of 0.0001, an Adam optimizer, hidden layers of size 124 and 24, a ReLU activation function, and was trained for 700 epochs. The number of epochs was based on exploratory work for a single random of fold of data, limited the risk of information leakage. The model had a total of 2,967,520 parameters.

Since many of the category/label combinations were missing, the missing labels were imputed before training to avoid having to apply masks. For every training fold, the missing outcome values were imputed in an iterative manner using a ridge regression model. Given the small sample size, a dedicated testing set was not possible, so instead performance was measured “out-of-fold”. A total of 5 folds were used during cross-validation.

## Results

To assess model performance, the out-of-fold predicted length-adjusted phenotypes were compared to their actual values. Figure 2 shows that the overall rank correlation between the predicted and actual length-adjusted outcomes ranged from 1-31%. Correlation across categories and labels was highest for Spearman’s rho (20%) and more conservative for the concordance approaches (14%). The average correlation over all label types and correlations approaches was highest for pancreatic insufficiency (20%) and the infection rate (18%), and less high for sweat chloride (8%).

**Fig. 2.**
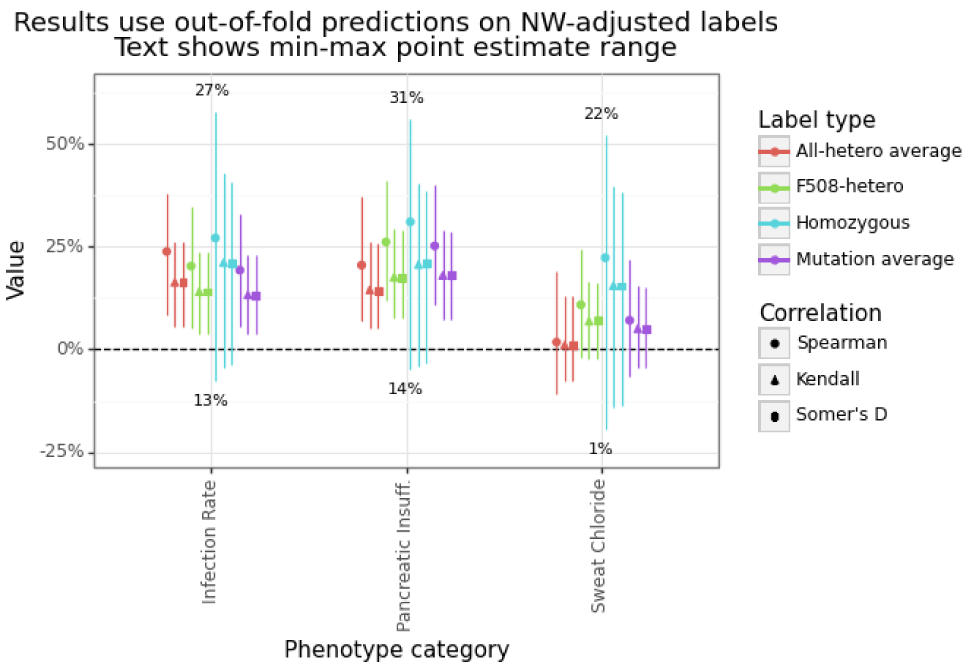
Predictive correlation for ESMFold-NNet model

The predicted to length-adjusted actual phenotype value can be seen in Figure S13 for all label types. Some labels tend to have more missing data (homozygous), while sweat chloride category has the most samples overall. The figure highlights the relative position of the predicted phenotypes for F508del and R347H, the former being a more severe mutation compared to the latter, with the predicted values reflect this difference.

To compare the increase in performance beyond knowing the amino acid sequence along, the NW kernel regression prediction was added to the ESMFold-NNet output (aka NW+NNET). Figure 3 shows that the NNet’s predicted phenotype value is able to increase performance for most categories and label types. Interestingly, the order of performance for the stacked model is reversed. Now the improvement for the Infection Rate category ranges from 16-22%, the Sweat Chloride category from (25 to 27%), and Pancreatic Insufficiency category from (32 to 44%). All results are statistically significant. Figure S14 shows the predicted vs actual outcome for both the NW and NW+NNet model. The NW predictions tend to be flat for most of the predicted range except for the CFTR mutations close to the wild-type length.

**Fig. 3.**
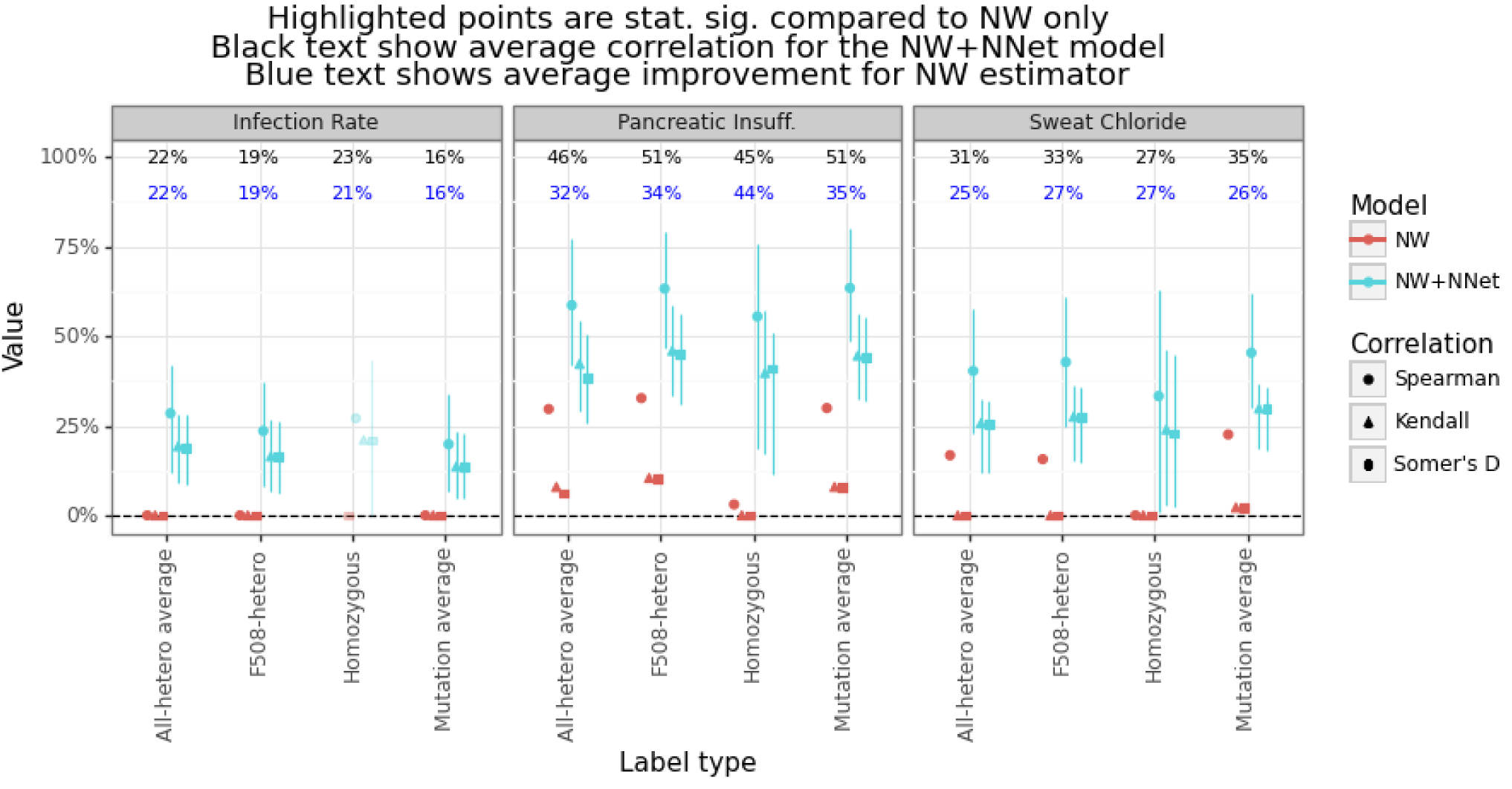
Improvement in correlation from ESMFold-NNet model (negative NW performance set to zero)

## Conclusions

This analysis has shown how a predictive ML algorithm can be trained predict the severity of CF-related phenotypes using embeddings from the ESMFold model. The model is able to demonstrate non-trivial and statistically significant performance for a variety of rank order measures for three phenotype categories (sweat chloride levels, infection rates, and pancreatic insufficiency) for multiple approaches. There does not appear to be any signal for predicting lung function.

There are many directions in which this research could be extended including: i) improvements to the processing pipeline to obtain more samples, ii) the use of non-censored clinical values from the CFTR2 database, iii) multiple forward passes through ESMFold (i.e. increasing the number of recycles), iv) experimenting with different regression model architectures, and v) weighting labels by sample size during training and evaluation.

Overall, linking CF phenotypes from the CFTR2 database to representations of core biological structures provides a new approach for classifying the pathogenicity of CFTR mutations in terms of clinical outcomes.

## ACKNOWLEDGEMENTS

Special thanks to Sunyun Lee for her research advice.

## Supplementary Note S1: Processing pipeline

The pipeline used to generate all data and results are schematically shown in Figure S1. A repo that contains all code to reproduce the analysis are found here: cftr2_esmfold. The rest of the section is as follows: Subsection A gives an overview of the clinical and genetic data, Subsection B describes how inferences was done with the ESMFold model, Subsection C gives additional information around the labels and features, and Section S2 contains all supplementary figures.

**Fig. S1.**
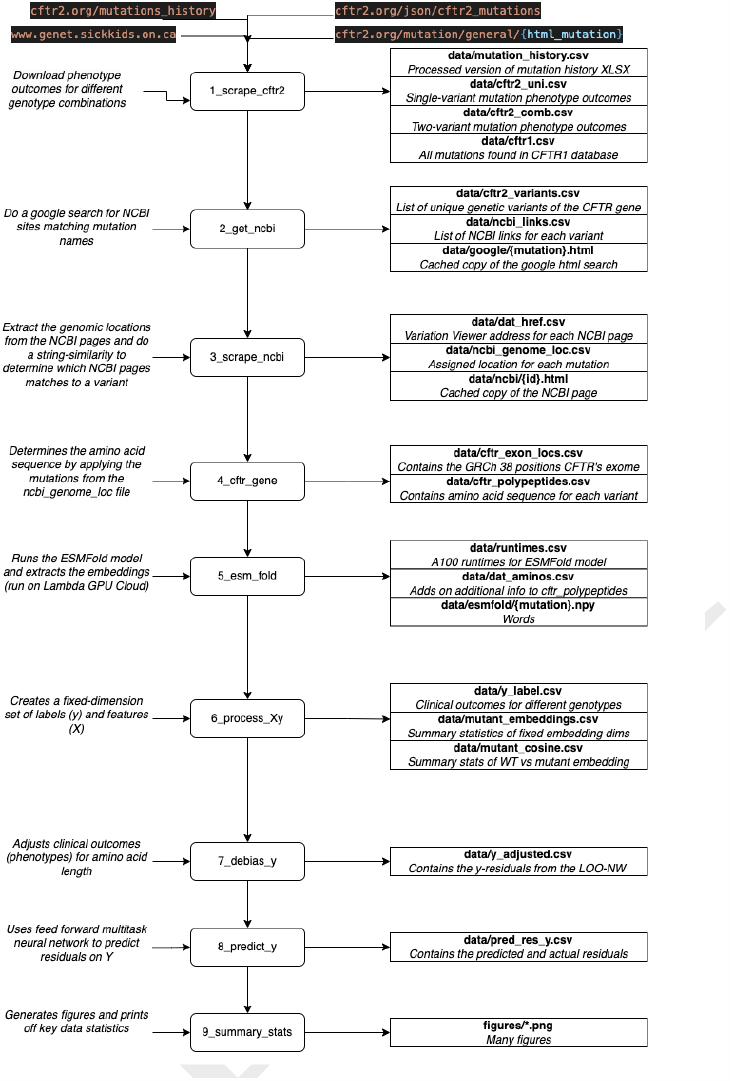
Processing pipeline overview of the cftr2_esmfold repo

### A. Clinical and variant data

The CFTR2 database imposes small-cell censoring, with any genotype that has 5 or fewer observation being masked. This means that if the data is missing (censored) for a single-variant, it missing for a two-variant combination since the population size can only shrink. A total of 417 single-variant mutation data were obtained.

Of these 417 mutations, 296 had at least one measurement for the single-variant case, and thus a total of 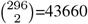 plus 296 homozygous pairs could possibly have data for two-variant pairs. Instead of running all of these combinations, the allele frequency of in the individual variants was multiplied together, and the top 25K of the sorted top two-variant pairs was used. Past the first thousand or so ranked pairs, the odds of finding one that has a CFTR2 page, let alone non-missing data becomes increasingly rare. A total of 1885 mutation pairs were obtained from CFTR2 (of which 294 are F508 heterozygous and 169 are homozygous).

Figure S2 provides a visual breakdown of these numbers, where “number of measurements” is the number of non-missing measurements for one of the four phenotypes, and “paired alleles” refers to whether it is the single or two-variant combination. The number of single-variant genotypes with at least one value (296) represents most of the data (70%), with most of these having three or four measurements (out of four). In contrast for the two-variant combinations, only 369 of the 1885 mutations (20%) hade at least measurement. Of these 369 non-completely missing two-variant pairs, F508del accounts for 219 of them (60%), and there were a total of 294 two-variant F508del pairs (75%). In summary, the two-variant pair is very much an F508del affair.

**Fig. S2.**
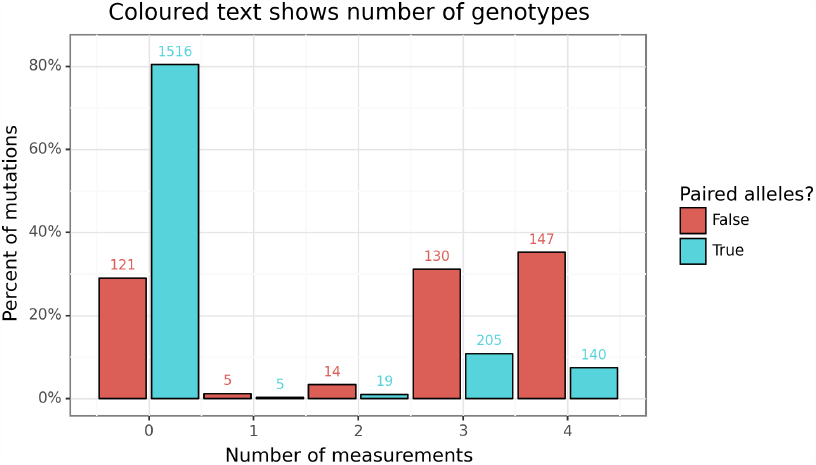
Number of mutations with data

For each of the 417 variants, an attempt was made to link them to an NCBI page through a simple google search. Any variants with matches (there could be more than one), had their genomic location extracted from the corresponding address of the GRCh38 NCBI Variation Viewer. A final match was determined by comparing the name found on the NCBI page to the original mutation name. The CFTR2 database had three names for mutations: a cDNA name, a protein name, and a legacy name. The closest string match (measured in Levenshtein distance) for each mutation is then selected. Any match that has a Levenshtein distance of 10 or greater was removed. If the cDNA and protein name align (in terms of genomic coordinates), then this value is chosen. For the mutations that disagree, cDNA match was chosen. The legacy name was not considered.

For each mutation, a simple transformation was done to the DNA sequence to determine the new sequence of amino acids associated with that protein. The reference DNA sequence was based on GRCh38 and Ensembl 109. The wild-type CFTR gene is made up 27 exons and 4443 base pairs (which includes a stop codon). Figure S8 shows the empirical CDF of the amino acid length for each version of mutated CFTR gene. The wild-type CFTR gene has 1480 amino acids, and around half of the exonic CFTR mutations are (approximately) this length. Mutant CFTR genes can have significantly fewer amino acids than the wild-type due to mutations which add a stop codon.

### B. ESMFold

As noted in the main body of text, two approaches were used to generate fixed dimensions from the ESMFold embeddings.

The first approach simply calculates the mean over all dimensions except the last. For example, s_s is a (1, n_amino, 1024) matrix, so we simply take the mean over dimensions 1 & 2 and get back a vector of length 1024. This is repeated for the min, max, and standard deviation. The final dimensionality is equivalent to 4(384 (states) + 1024 (s_s) + 128 (s_z))=6144.

The second approach is a bit more involved. Before calculating the cosine similarity, the matrices are reduced to two dimensions. For s_s this is easy, and the first dimension is simply flattened ((1, n_amino, 1024) becomes (n_amino, 1024)). For states, the average is taken ((8,1,n_amino,384) becomes (n_amino,384)). For s_z the diagonal is taken ((1, n_amino, n_amino, 128) becomes (n_amino, 128)). Next, we calculate the cosine similarity between the wild-type and the mutant for each, and get back three matrices of dimension: (1480,n_amino), where the i,j’th entry corresponds to the cosine similarity of wild-type dimension i and mutant dimension j for the respective embeddings. Like the first approach, the mean, min, max, and standard deviation are calculated across the rows giving a total of 4 ·3 · 1480 = 17760 features. These features can be concatenated to obtain a final feature vector of length 23,904 for each mutation.

During inference, 311 amino acid sequences were passed through the model. Each mutant amino acid sequence was recorded until the first stop codon. Amino acids sequences that were less than a length of 100 were excluded. The following parameters were set:

1. num_recycles=1: The forward pass was only run done once for each mutation. However, this number could likely be increased for better results, as is often found in the literature for protein folding problem.
2. chain_linker=25: Represents the number of residues in the linker between chains
3. chunk_size=64: Represents that size of the cache used doing the forward pass

### C. Modelling

A total of 225 samples were available for training and evaluation. This means 260 CFTR mutants were removed during the processing pipeline, for which Figure S3 provides a breakdown. Of these 260, 105 were due to external factors (mutations not occurring on the exonic regions (31) or lacking data in CFTR2 due to small-cell censoring (74)), 107 were due to data collection issues (not all URLs were parsed (68) and not all mutants could find an NCBI page (39)), and 48 were due to processing choices (short protein sequences were removed (38), along with missense (2) and duplicate (8) mutants). Of these data losses, only the exonic regions is structural, suggesting future work could help to double the sample size by adding another 229 mutations.

**Fig. S3.**
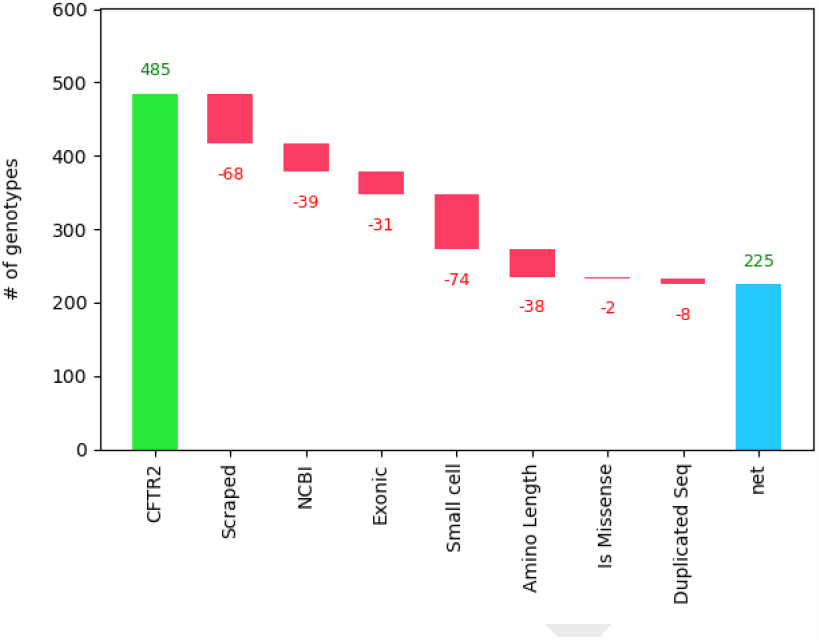
Factors determining final dataset size

Early exploratory work revealed that predictive models for the lung function categories (for all label types) showed performance largely no better than random guessing (see [this figure](https://github.com/ErikinBC/cftr2_esmfold/blob/main/figures/oof_perf_lung.png)). The final model run used only 12 labels from the combination of three categories (sweat chloride, pancreatic insufficiency, and infection rate). A simple two-layer neural network model using the [MLPRegressor](https://scikit-learn.org/stable/modules/generated/sklearn.neural_network.MLPRegressor.html) model class was used to predict all 12 outcomes simultaneously. The NNet used a learning rate of 0.0001, an Adam optimizer, hidden layers of size 124 and 24, a ReLU activation function, and was trained for 700 epochs. The number of epochs was based on exploratory work for a single random of fold of data, limited the risk of information leakage. The model had a total of 2,967,520 parameters.

Since many of the category/label combinations were missing, I opted to impute the missing labels before training to avoid having to apply masks. During inference time this was not an issue since only the actual values could be compared to the predictions. For every training fold, the missing outcome values were imputed in an iterative manner using the [IterativeImputer](https://scikit-learn.org/stable/modules/generated/sklearn.impute.IterativeImputer.html) class with the [BayesianRidge](https://scikit-learn.org/stable/modules/generated/sklearn.linear_model.BayesianRidge.html) regression model.

## Supplementary Note S2: Figures

**Fig. S4.**
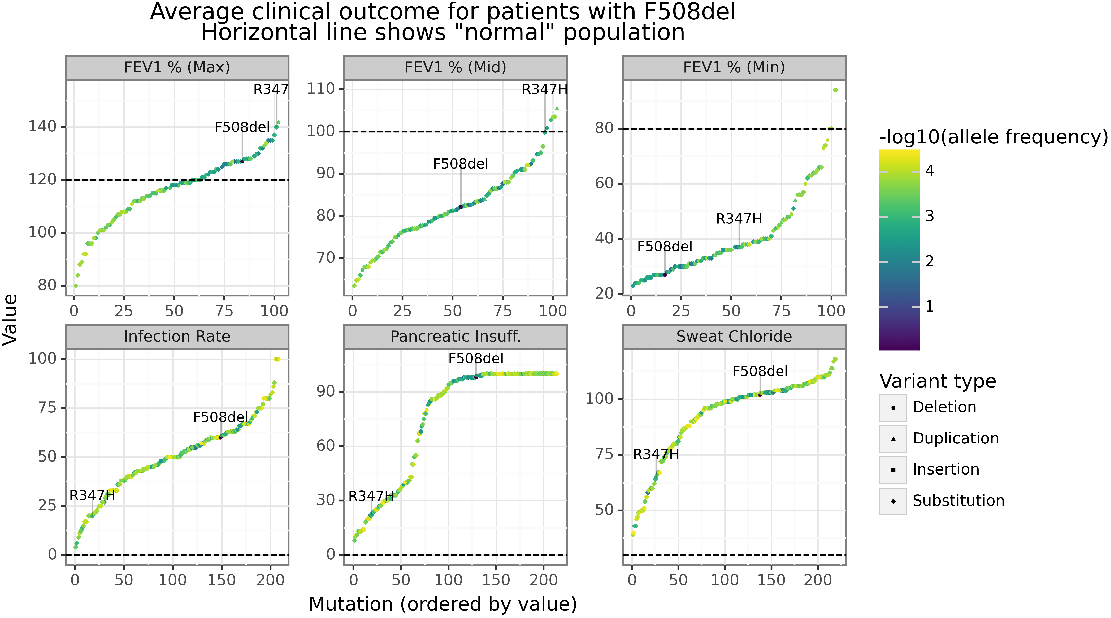
Distribution of average clinical outcomes across 100+ CFTR2 genotypes

**Fig. S5.**
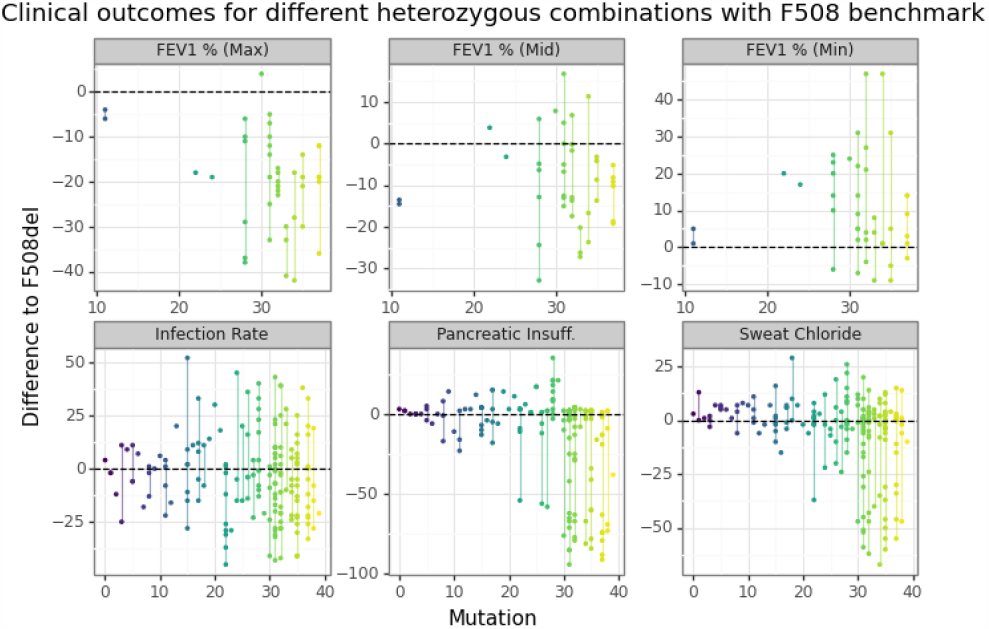
Range of heterozygous outcomes for F508del

**Fig. S6.**
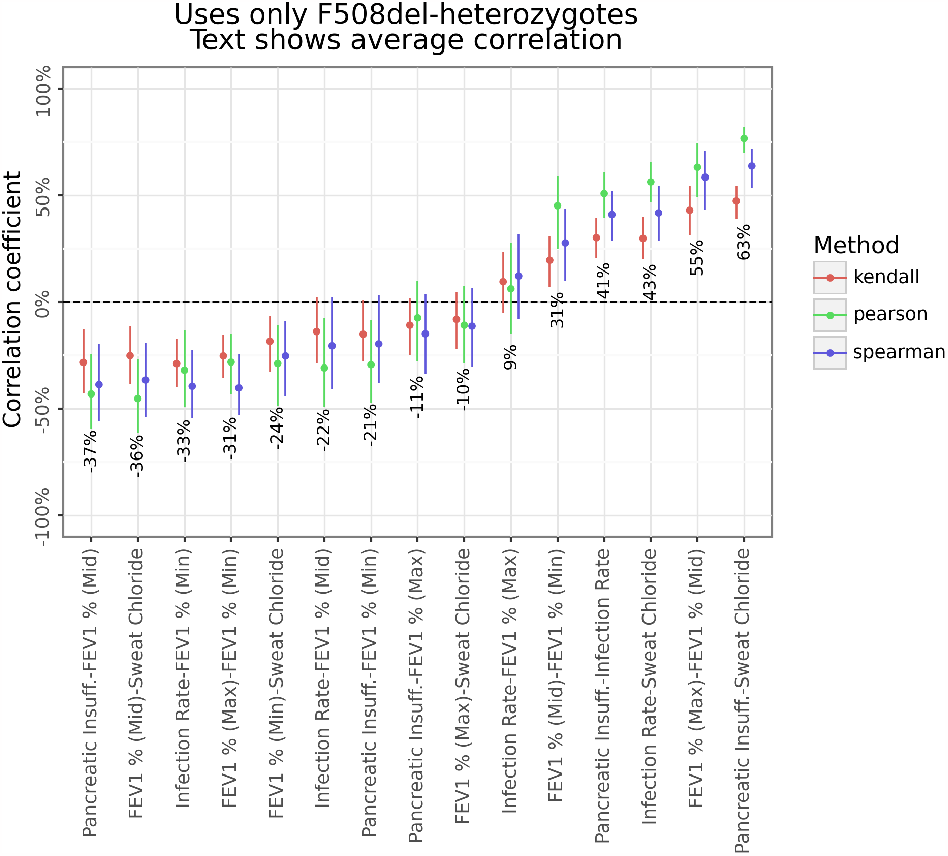
Between category correlation

**Fig. S7.**
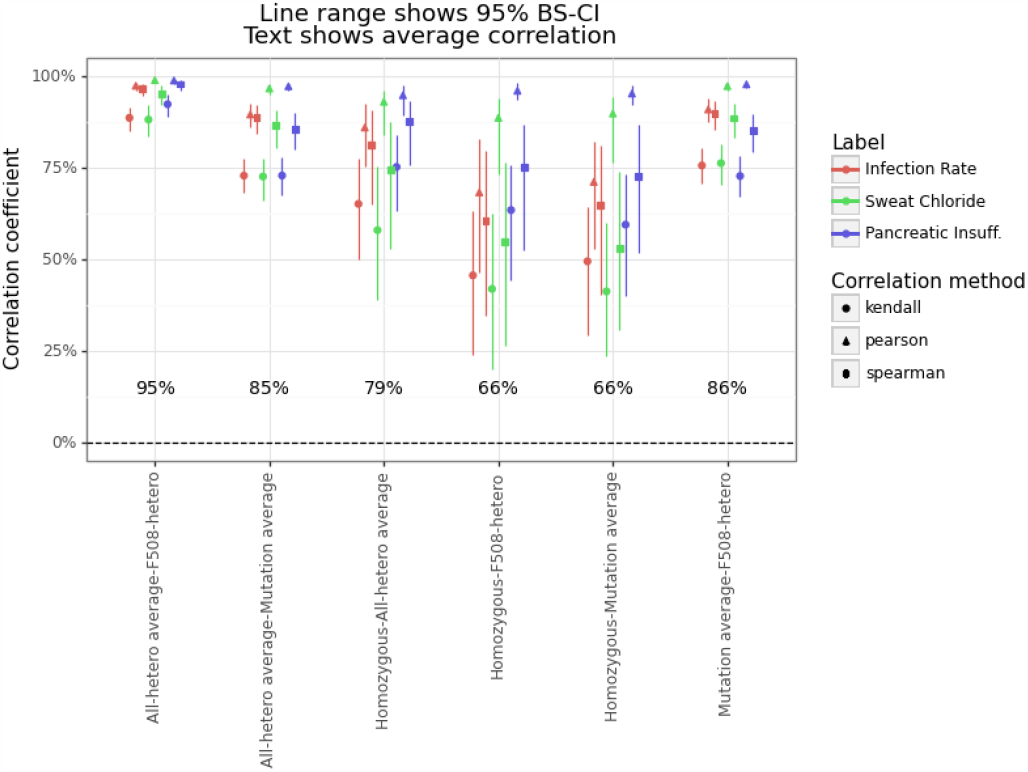
Within category correlation

**Fig. S8.**
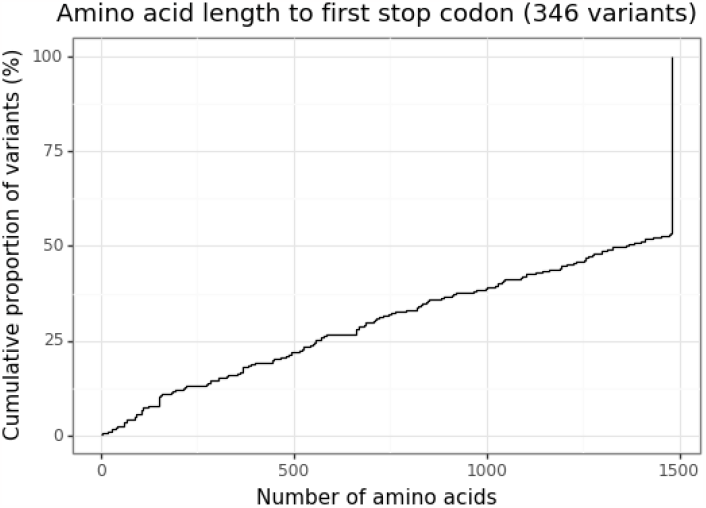
Mutant CFTR gene lengths to first stop codon

**Fig. S9.**
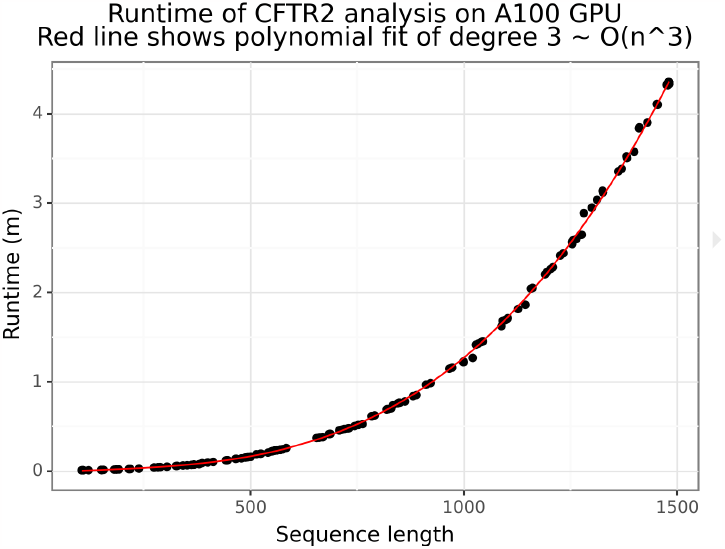
ESMFold has a polynomial runtime

**Fig. S10.**
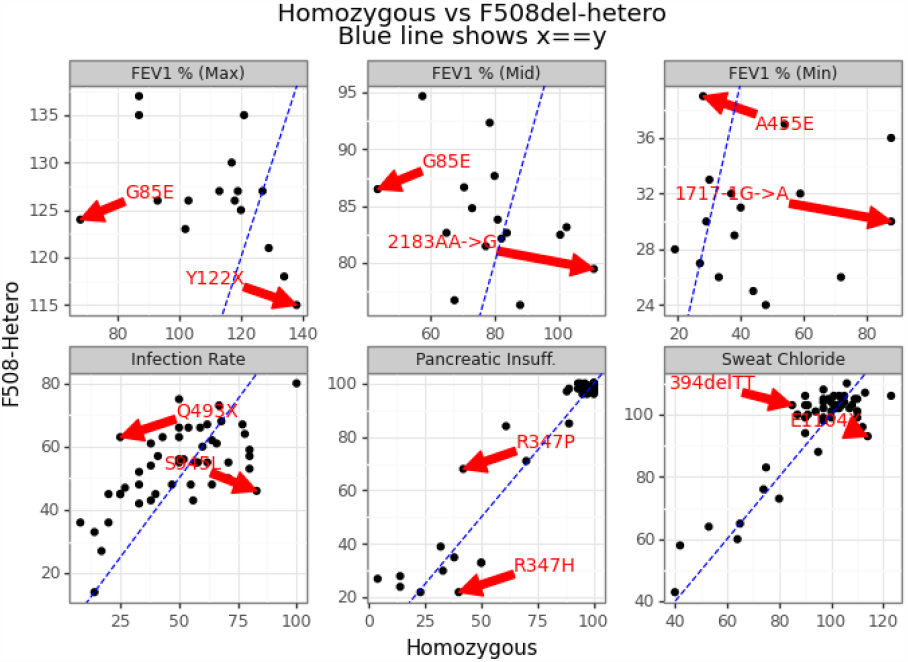
Relationship between homozygous and F508del-heterozygous

**Fig. S11.**
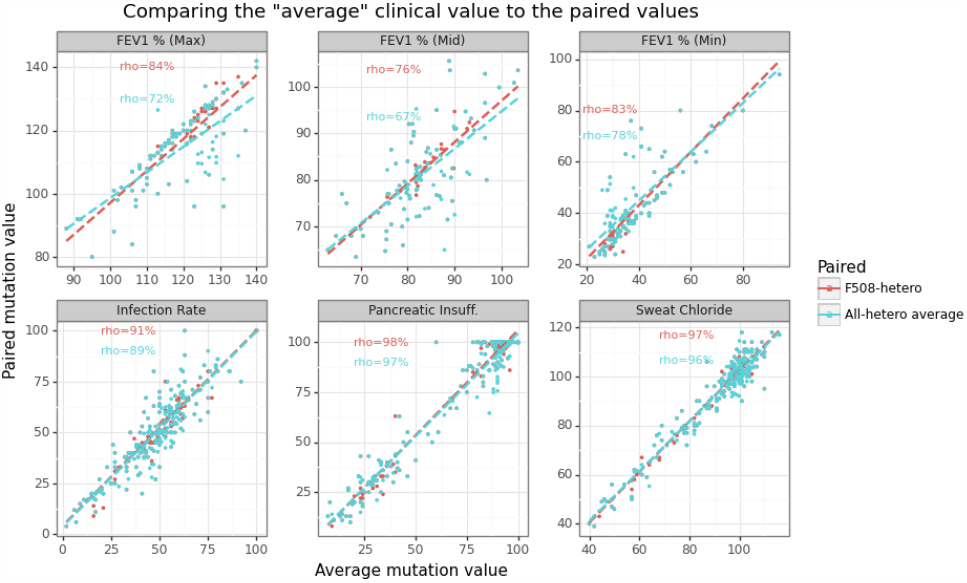
Relationship between allele average and heterozygotes

**Fig. S12.**
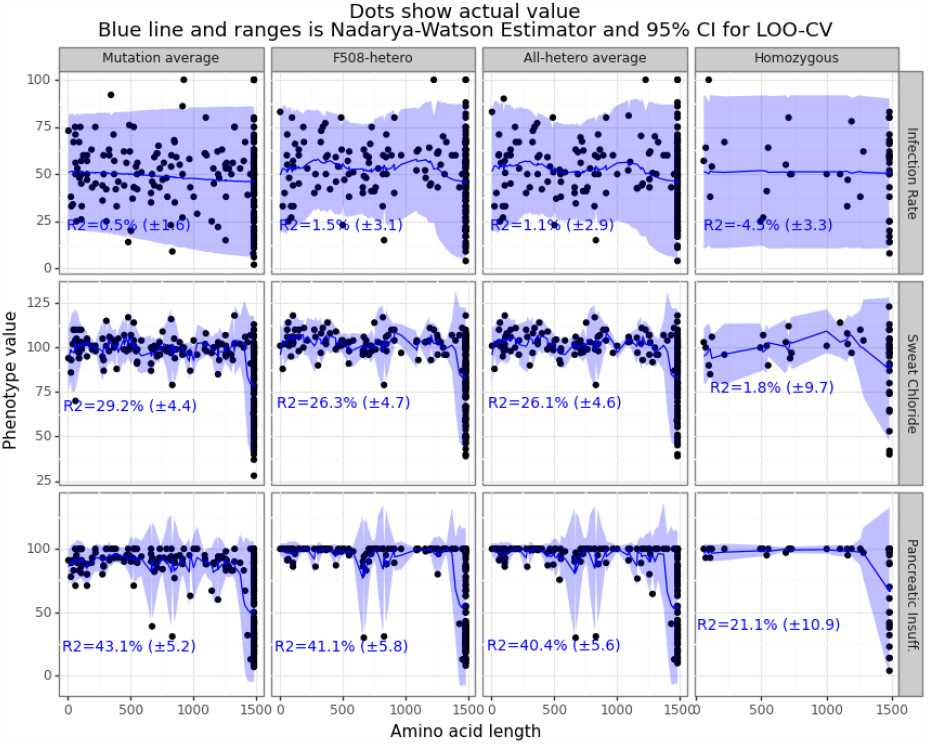
N-W kernel regression adjusts clinical outcomes

**Fig. S13.**
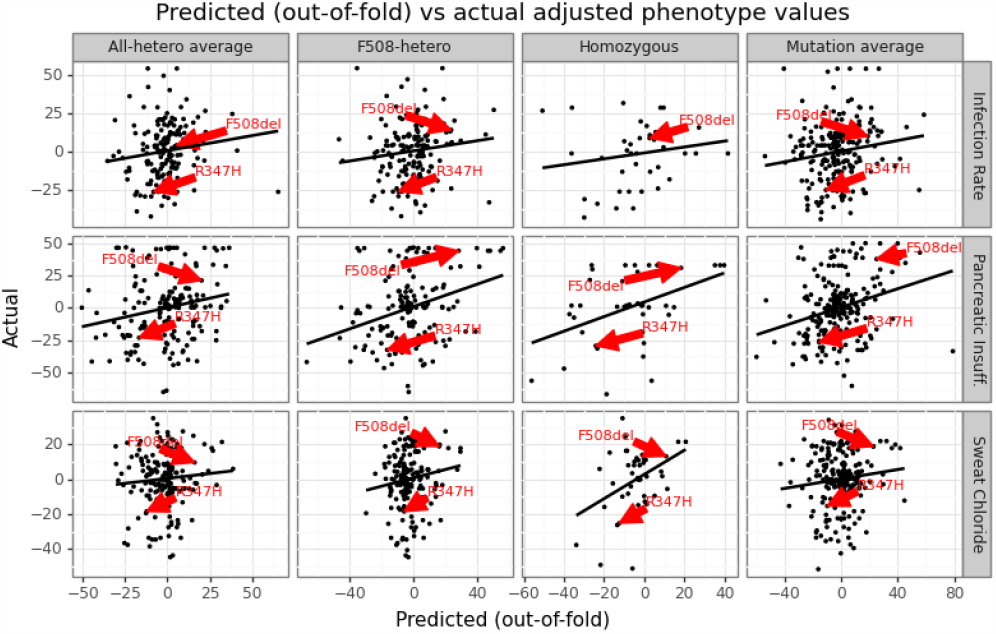
Predicted vs actual (adjusted outcome) for ESMFold-NNet model

**Fig. S14.**
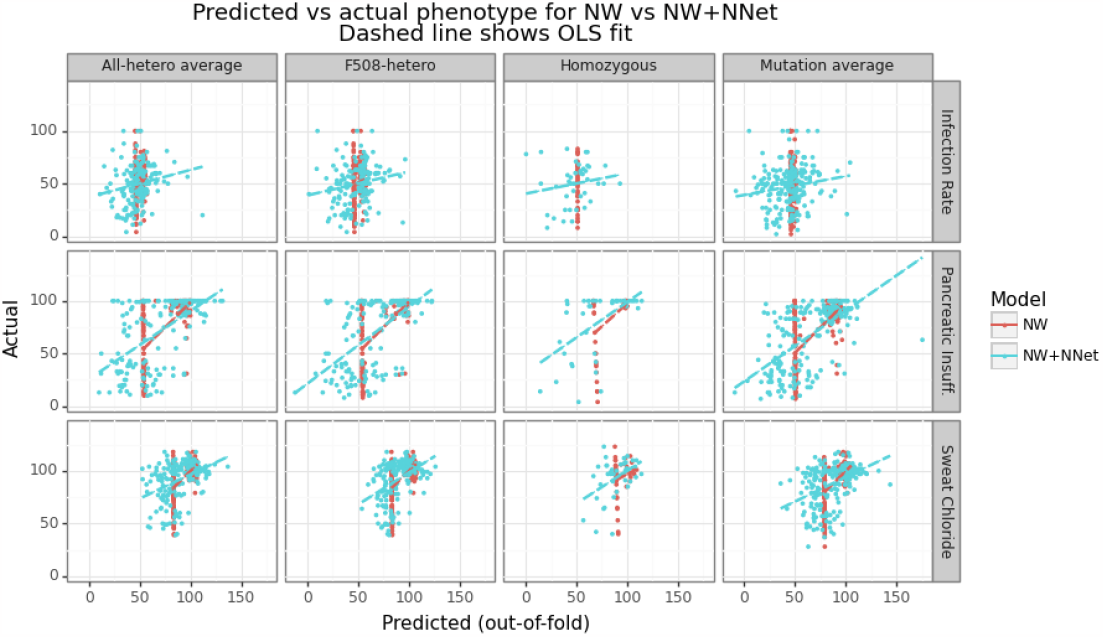
Predicted vs actual for ESMFold-NNet model

## Footnotes

1 See model documentation for more information.

## Notes

https://github.com/ErikinBC/cftr2_esmfold

### Competing Interest Statement

The authors have declared no competing interest.

### Summary of Updates

Fixing duplicate sentences in intro section

