## Supplementary figures and tables for "A multitask neural network trained on embeddings from ESMFold can accurately rank order clinical outcomes for different cystic fibrosis mutations"

### Supplementary Note S1: Processing pipeline

The pipeline used to generate all data and results are schematically shown in Figure S1. A repo that contains all code to reproduce the analysis are found here: cftr2_esmfold. The rest of the section is as follows: Subsection A gives an overview of the clinical and genetic data, Subsection B describes how inferences was done with the ESMFold model, Subsection C gives additional information around the labels and features, and Section S2 contains all supplementary figures.

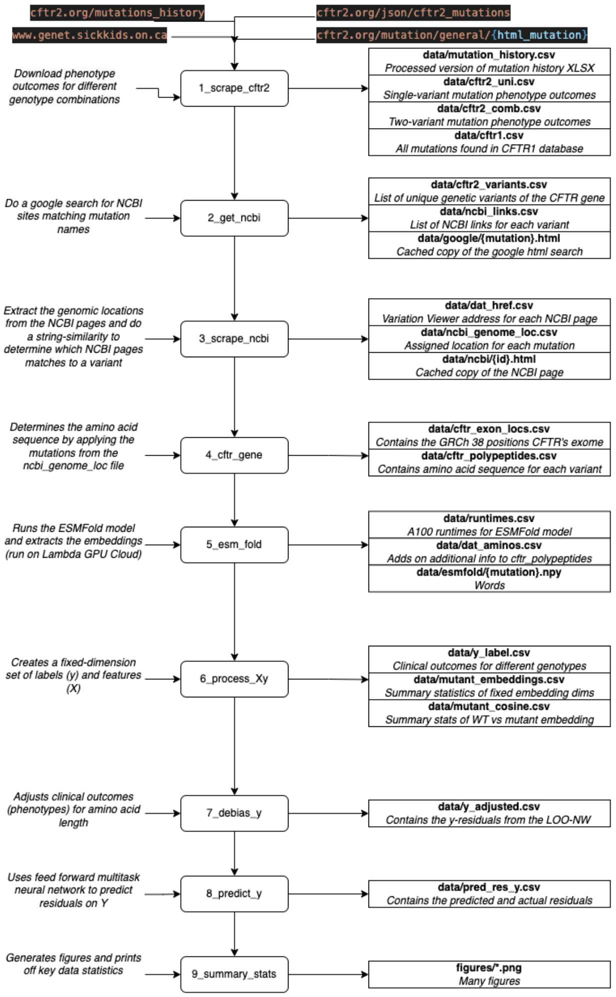


**Fig. S1.** Processing pipeline overview of the cftr2_esmfold repo

**A. Clinical and variant data.** The CFTR2 database imposes small-cell censoring, with any genotype that has 5 or fewer observation being masked. This means that if the data is missing (censored) for a single-variant, it missing for a two-variant combination since the population size can only shrink. A total of 417 single-variant mutation data were obtained.

Of these 417 mutations, 296 had at least one measurement for the single-variant case, and thus a total of $\binom{296}{2}$=43660 plus 296 homozygous pairs could possibly have data for two-variant pairs. Instead of running all of these combinations, the allele frequency of in the individual variants was multiplied together, and the top 25K of the sorted top two-variant pairs was used. Past the first thousand or so ranked pairs, the odds of finding one that has a CFTR2 page, let alone non-missing data becomes increasingly rare. A total of 1885 mutation pairs were obtained from CFTR2 (of which 294 are F508 heterozygous and 169 are homozygous).

Figure S2 provides a visual breakdown of these numbers, where "number of measurements" is the number of non-missing measurements for one of the four phenotypes, and "paired alleles" refers to whether it is the single or two-variant combination. The number of single-variant genotypes with at least one value (296) represents most of the data (70%), with most of these having three or four measurements (out of four). In contrast for the two-variant combinations, only 369 of the 1885 mutations (20%) hade at least measurement. Of these 369 non-completely missing two-variant pairs, F508del accounts for 219 of them (60%), and there were a total of 294 two-variant F508del pairs (75%). In summary, the two-variant pair is very much an F508del affair.

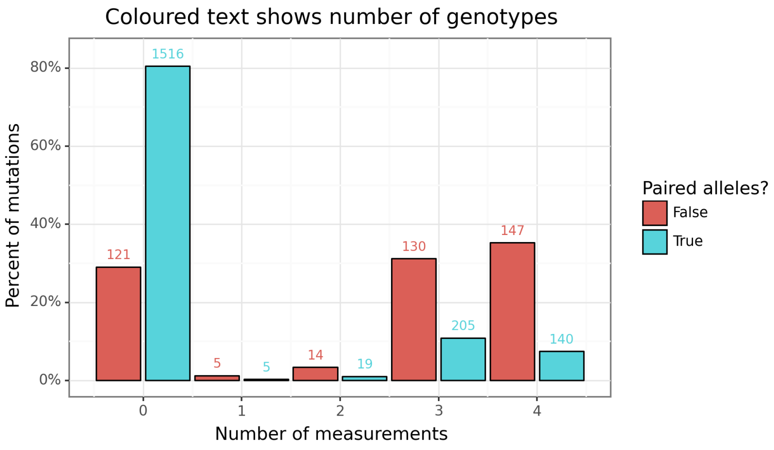


**Fig. S2.** Number of mutations with data

For each of the 417 variants, an attempt was made to link them to an NCBI page through a simple google search. Any variants with matches (there could be more than one), had their genomic location extracted from the corresponding address of the GRCh38 NCBI Variation Viewer. A final match was determined by comparing the name found on the NCBI page to the original mutation name. The CFTR2 database had three names for mutations: a cDNA name, a protein name, and a legacy name. The closest string match (measured in Levenshtein distance) for each mutation is then selected. Any match that has a Levenshtein distance of 10 or greater was removed. If the cDNA and protein name align (in terms of genomic coordinates), then this value is chosen. For the mutations that disagree, cDNA match was chosen. The legacy name was not considered.
For each mutation, a simple transformation was done to the DNA sequence to determine the new sequence of amino acids associated with that protein. The reference DNA sequence was based on GRCh38 and Ensembl 109. The wild-type CFTR gene is made up 27 exons and 4443 base pairs (which includes a stop codon). Figure S8 shows the empirical CDF of the amino acid length for each version of mutated CFTR gene. The wild-type CFTR gene has 1480 amino acids, and around half of the exonic CFTR mutations are (approximately) this length. Mutant CFTR genes can have significantly fewer amino acids than the wild-type due to mutations which add a stop codon.

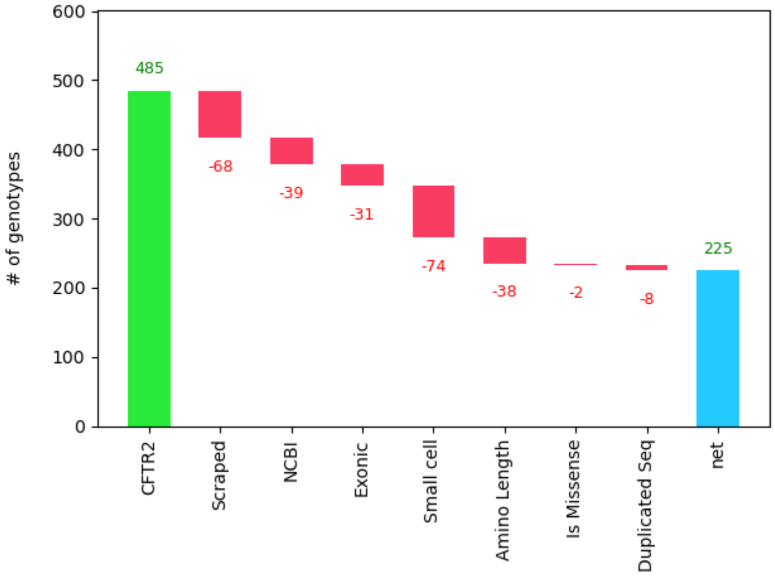


**Fig. S3.** Factors determining final dataset size

Early exploratory work revealed that predictive models for the lung function categories (for all label types) showed performance largely no better than random guessing (see this figure). The final model run used only 12 labels from the combination of three categories (sweat chloride, pancreatic insufficiency, and infection rate). A simple two-layer neural network model using the MLPRegressor model class was used to predict all 12 outcomes simultaneously. The NNet used a learning rate of 0.0001, an Adam optimizer, hidden layers of size 124 and 24, a ReLU activation function, and was trained for 700 epochs. The number of epochs was based on exploratory work for a single random of fold of data, limited the risk of information leakage. The model had a total of 2,967,520 parameters.

Since many of the category/label combinations were missing, I opted to impute the missing labels before training to avoid having to apply masks. During inference time this was not an issue since only the actual values could be compared to the predictions. For every training fold, the missing outcome values were imputed in an iterative manner using the IterativeImputer class with the BayesianRidge regression model.

DRAFT

### Supplementary Note S2: Figures

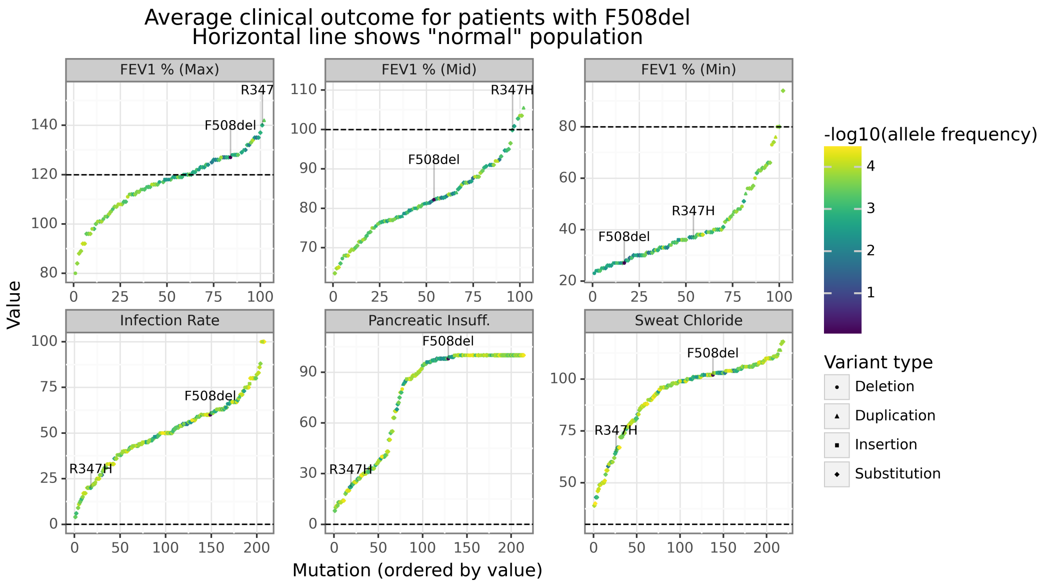


**Fig. S4.** Distribution of average clinical outcomes across 100+ CFTR2 genotypes

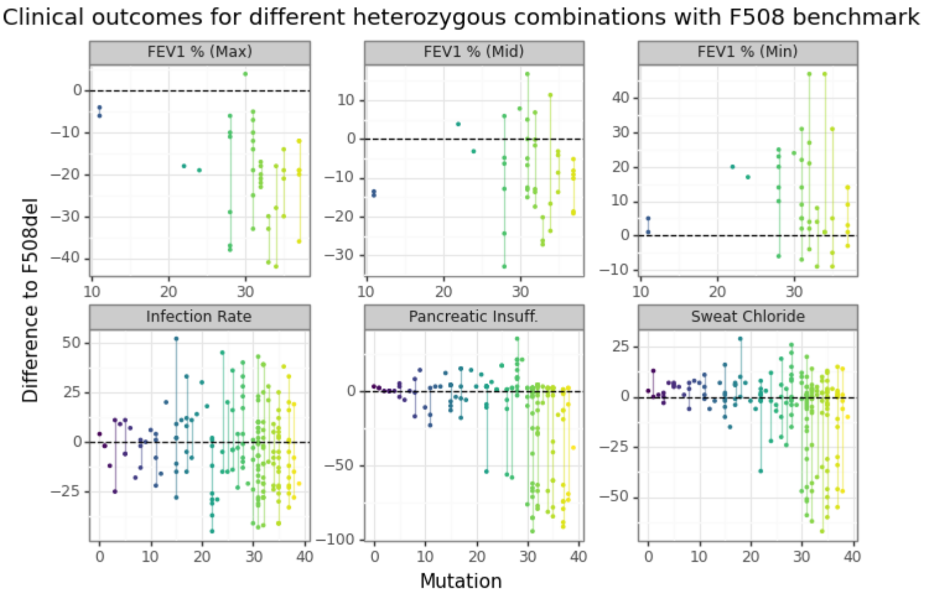


**Fig. S5.** Range of heterozygous outcomes for F508del

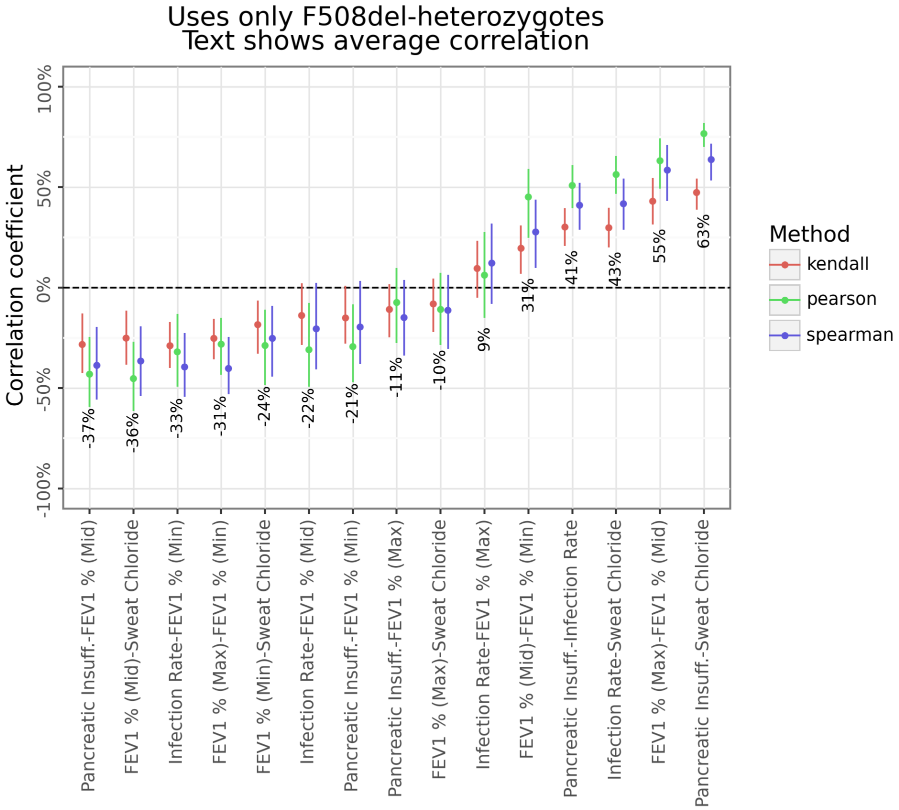


**Fig. S6.** Between category correlation

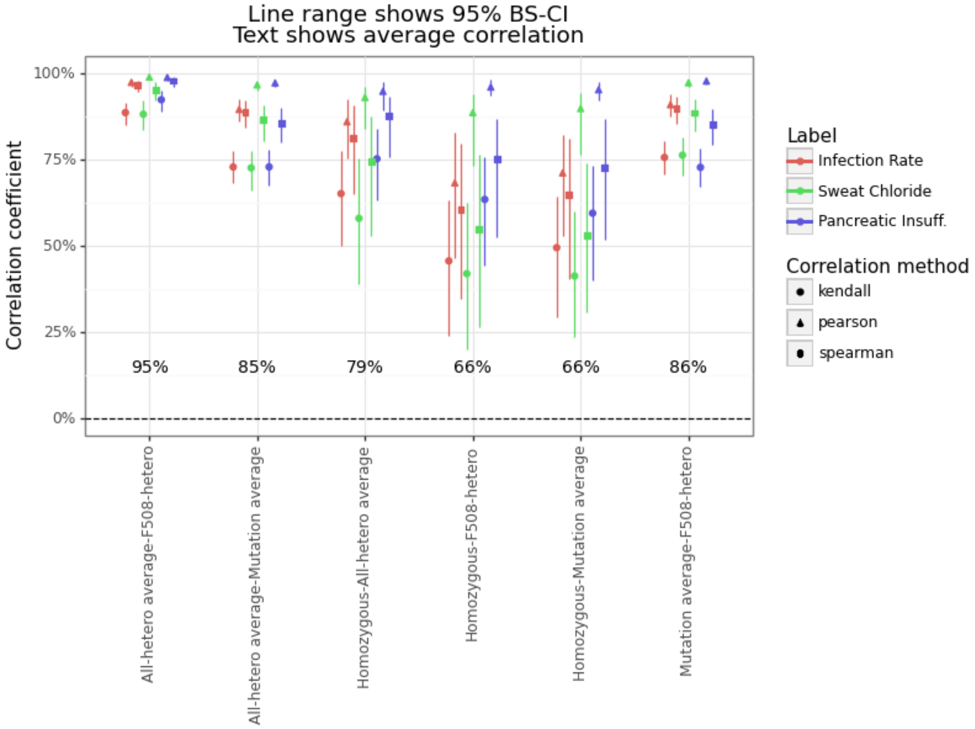


**Fig. S7.** Within category correlation

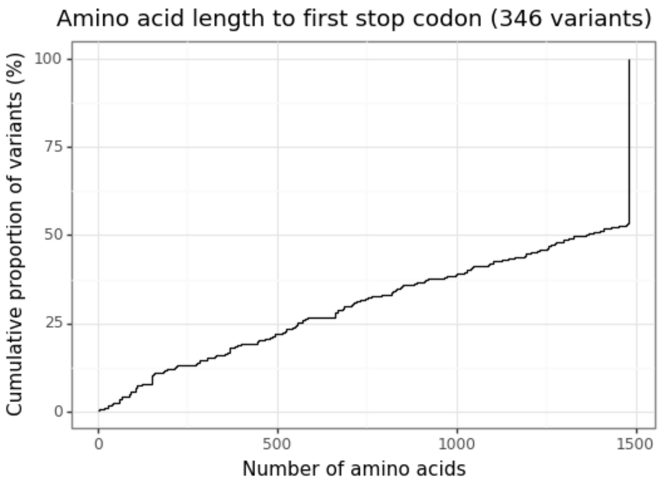


**Fig. S8.** Mutant CFTR gene lengths to first stop codon

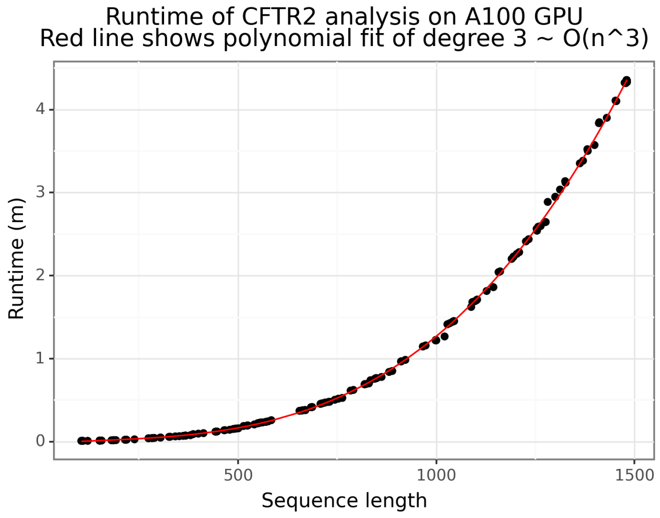


**Fig. S9.** ESMFold has a polynomial runtime

Dr

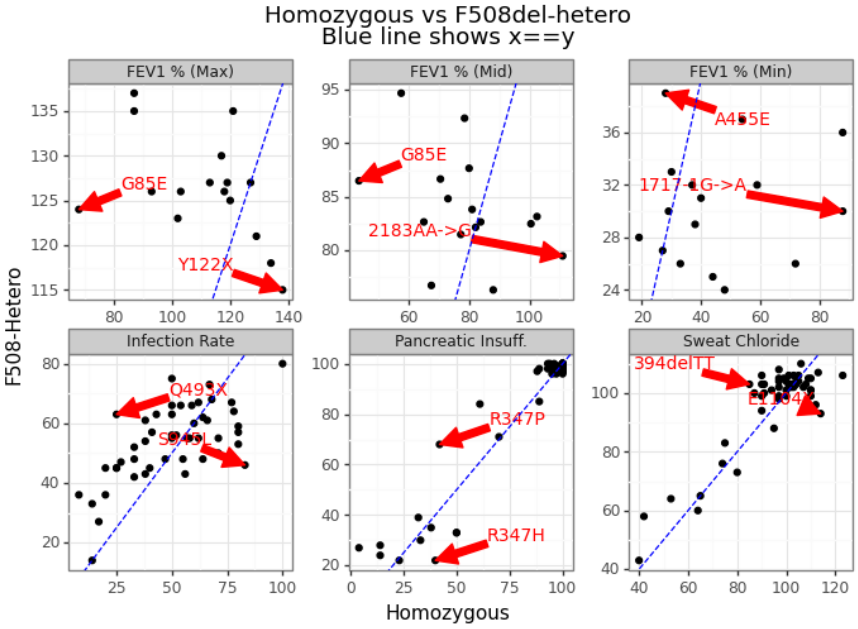


**Fig. S10.** Relationship between homozygous and F508del-heterozygous

DRAFT

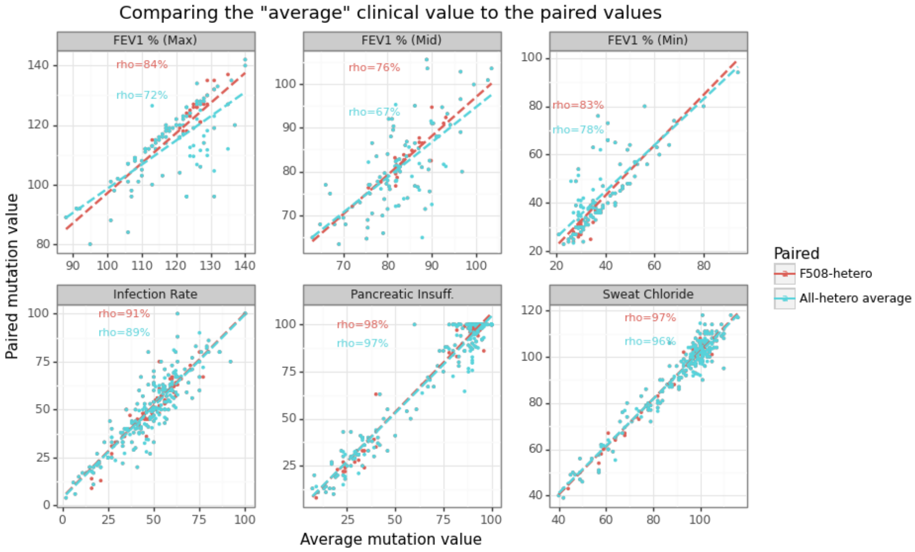


**Fig. S11.** Relationship between allele average and heterozygotes

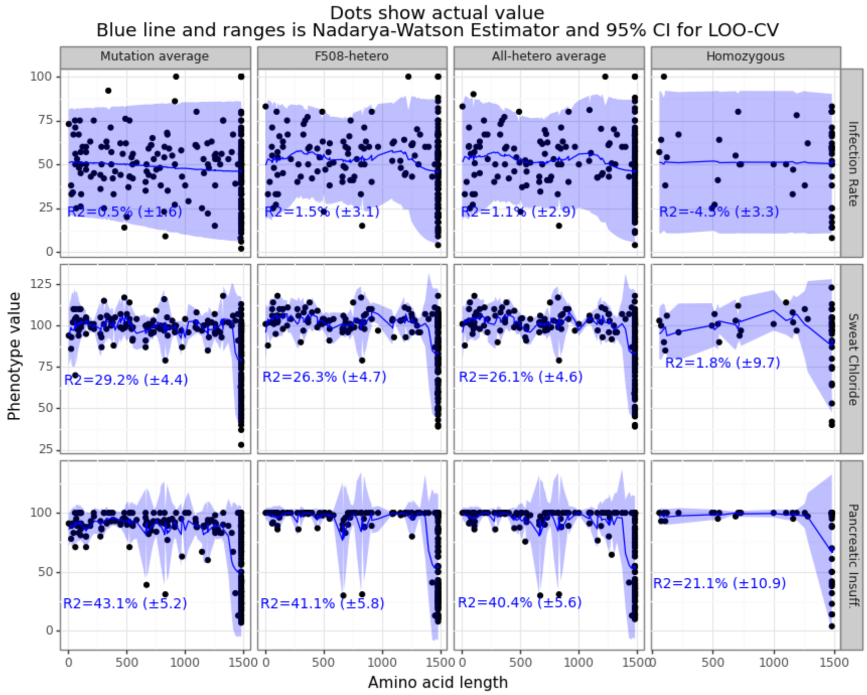


**Fig. S12.** N-W kernel regression adjusts clinical outcomes

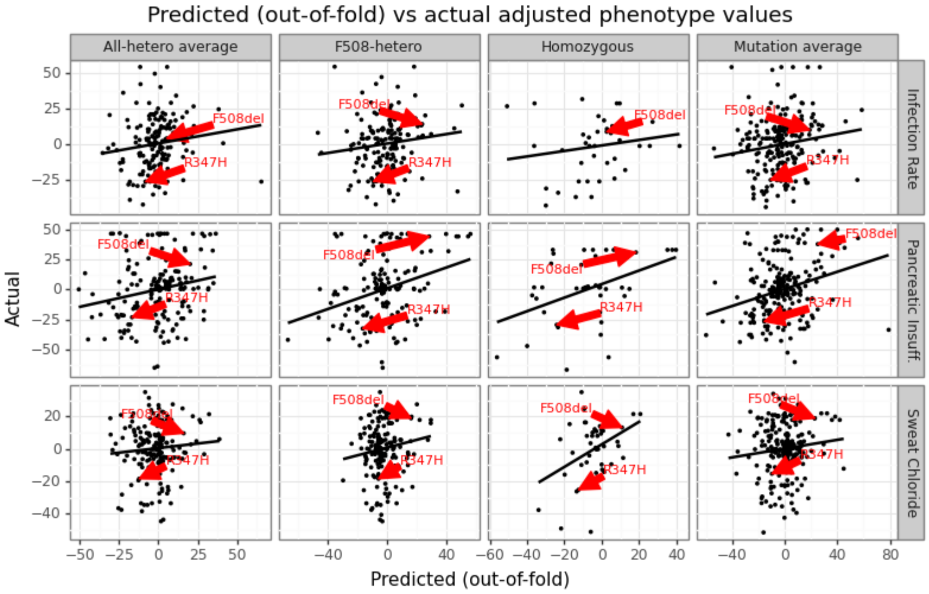


**Fig. S13.** Predicted vs actual (adjusted outcome) for ESMFold-NNet model

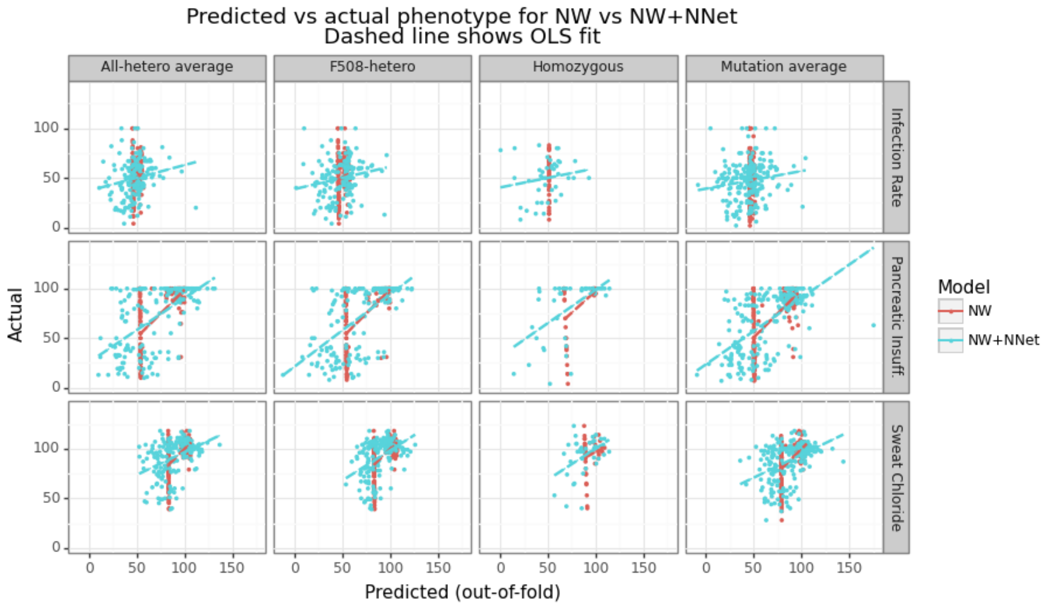


**Fig. S14.** Predicted vs actual for ESMFold-NNet model
